# Improved Diagnostic of Spotted Fevers via Loop-Mediated Isothermal Amplification with One Step Strand Displacement

**DOI:** 10.1101/2023.02.07.526334

**Authors:** Simren Lakhotia, Timothy E. Riedel

## Abstract

Spotted fever rickettsiosis plagues countries around the world. One of the deadliest of this group, Rickettsia *rickettsii*, responsible for Rocky Mountain spotted fever, is an emerging tickborne illness in North America. The predominant clinical diagnostic is PCR based but does not work until disease has progressed to a severe phase of infection, at which point the outcome of a full recovery is significantly decreased. An alternative, loop mediated isothermal amplification through one-step strand displacement (LAMP-OSD) assay, was developed to improve diagnostic speed and sensitivity. Synthetic dsDNA genes from the 17 kDa surface antigen precursor (AY281069.1) amplified between fifteen minutes to an hour and were detected to concentrations as low as 10^2^ copies/μL. This RMSF LAMP-OSD assay shows promise to deliver results in just a few hours and the detection limit is potentially 100 times more sensitive than qPCR based assays.

## 2. Introduction

Tick and flea-borne rickettsia infections are often underfunded and an under-researched topic, especially in low-resource countries (Noden et al. 2018). One of the primary reasons these diseases go undetected is due to their low rickettsiae count during the early stages of infection, which current diagnostic measures are unable to pick up (Richards 2012). Rocky Mountain spotted fever in North and Central America, Mediterranean spotted fever and Israeli spotted fever in Europe, and African tick bite fever in Africa are some of the common spotted fevers. In the case of Far-Eastern spotted fever and Siberian tick typhus, both are tick-borne and present similarly, but one is a spotted fever while the other is a typhus, and thus are treated with different medications. These two diseases can only be distinguished through molecular diagnostic methods. All of these spotted fevers are a group of diseases caused by closely related bacteria (Parola et al. 2013, Phan et al. 2011), and are treated with doxycycline (Biggs et al. 2016, Agahan et al. 2011, Raoult et al. 2001). These bacteria are spread to people through the bite of an infected tick. The signs and symptoms of spotted fevers are nonspecific at first and can include: fever, headache, rash, nausea, vomiting, stomach pain, and muscle pain (Biggs et al. 2016, Agahan et al. 2001, Raoult et al. 2001). While many patients will develop tell-tale signs of the specific strain, be it a rash, ophthalmic disease, or eschars, these symptoms do not manifest soon enough to be used as the basis for diagnoses (Biggs et al. 2016, Raoult et al. 2001).

The main forms of lab confirmation of infection as it stands are serology and Polymerase Chain Reaction (PCR). Serology diagnosis is typically confirmed by documenting a four-fold or greater rise in antibody titer between acute- and convalescent-phase serum samples, where the convalescent-phase is usually two to four weeks after the illness has resolved. Due to the nature of this test, it is confirmation of infection more than a diagnostic, considering the patient needs to wait almost a month after the onset of the illness. As for PCR, spotted fevers do not circulate in large numbers in the blood until the disease has progressed to a severe phase of infection, at which point the outcome of a full recovery is significantly decreased (Biggs et al. 2016). Before, the chances of a false negative are quite high.

Due to the short diagnosis window and nonspecific symptoms, spotted fevers make a good candidate for developing a highly sensitive point-of-care diagnostic. Physicians recommend that patients bring in the tick that bit them (Mayo Clinic 2019, WebMD 2020)., which can allow for further testing on the tick. A recent study successfully identified various Rickettsia species in field-collected ticks and fleas using loop-mediated isothermal amplification (LAMP) assays (Noden et al., 2018). This study aims to build upon the Noden et al. 2018 system by creating a toe-hold based amplification reporter for the LAMP assay that fluoresces (Jiang et al. 2015). This fluorescence output should be easier to discriminate thus greatly increasing the assay sensitivity and the OSD reporter output offers a second error checking mechanism since only the correct amplification sequence will be reported.

## 3. Methods

### 3.1. LAMP-OSD Probe Design

A previously published LAMP reaction (Noden et al.) was modified by substituting the Loop B primer with a toe-hold strand exchange probe using the suggested guidelines (Jiang et al. 2015). Benchling was used to design a fluorophore-quencher duplex, where the fluorophore strand binds to a region between B1/B2 primers. The duplex is developed so that it is non-complementary to itself, preventing internal loop structures, with the fluorophore’s strand GC ratio of 47.6% (Tm 67.9 °C) and the quencher’s strand GC ratio 43.75% (Tm 62.6 °C). After a probe was designed based on the *R. rickettsii* strand, it was then cross checked with similar Rickettsia diseases using BLAST to verify its effectiveness on similar diseases (S1-S3). The probe design was tested in Nupack using the following parameters: 65°C, two strand species with complex size of two and salts at 0.05 M Na+ and 0.004 Mg2+. Testing showed that at a 1:1 ratio, 99.2% of the probe and quencher strands would form a hemiduplex with a secondary structure free energy of -18.91 kcal/mol. The fluorophore strand by themselves had a secondary structure free energy of 0.0 kcal/mol and was shown to not create any self-hybrizing loop structures. This ensures that the strand would bind to the target DNA rather than to itself.

### 3.2. Annealing the OSD Probe

The fluorophore and quencher labeled DNA strands for the probe construct (section 3.1) were ordered from (IDT, USA) with HPLC purification. Strands were resuspended in 1x Isothermal Buffer (contains 20mM Tris-HCl, 10mM (NH_4_)_2_SO_4_, 50mM KCl, 2mM MgSO_4_, 0.1% Tween 20) in two separate mixtures of 1:5 and 1:4 fluorophore:quencher ratio. Before use, the mixed fluorophore and quencher strands were annealed into hemiduplex probes by heating the mixtures to 95°C for 5 minutes and then allowing them to cool on a lab bench in a tube rack.

### 3.3. LAMP-OSD Reaction Setting

Each LAMP-OSD reaction totaled 25 μl using (New England Biolabs, Ipswitch, MA, USA unless otherwise stated) consisting of 3.0 μl nuclease free water, 2.5 μl 10x Isothermal Amplification Buffer, 2.5 μl 10 mM dNTP Solution Mix, 0.5 μl 100 mM MgSO4, 5.0 μl 5M Betaine (Sigma-Aldrich), and 2 μl 10x primer mix (5μM F3 primer, 5 μM B3 primer, 20μM FIP, 20 μM BIP, and 10 μM LoopF) (Table 1)(Noden et al. 2018), 2.0 μl BST 2.0 WarmStart DNA Polymerase (8 units/μl), 2.5 μl OSD Probe, and 1 μl of sample DNA. LAMP amplification reactions were heated at 65° C for an hour and a half in either a LightCycler 96 (Roche, NC, U.S.A.) or a heat block. Samples were created in white PCR tubes if run in the LightCycler 96, and clear PCR tubes if run in the heat block.

**Table 1:**
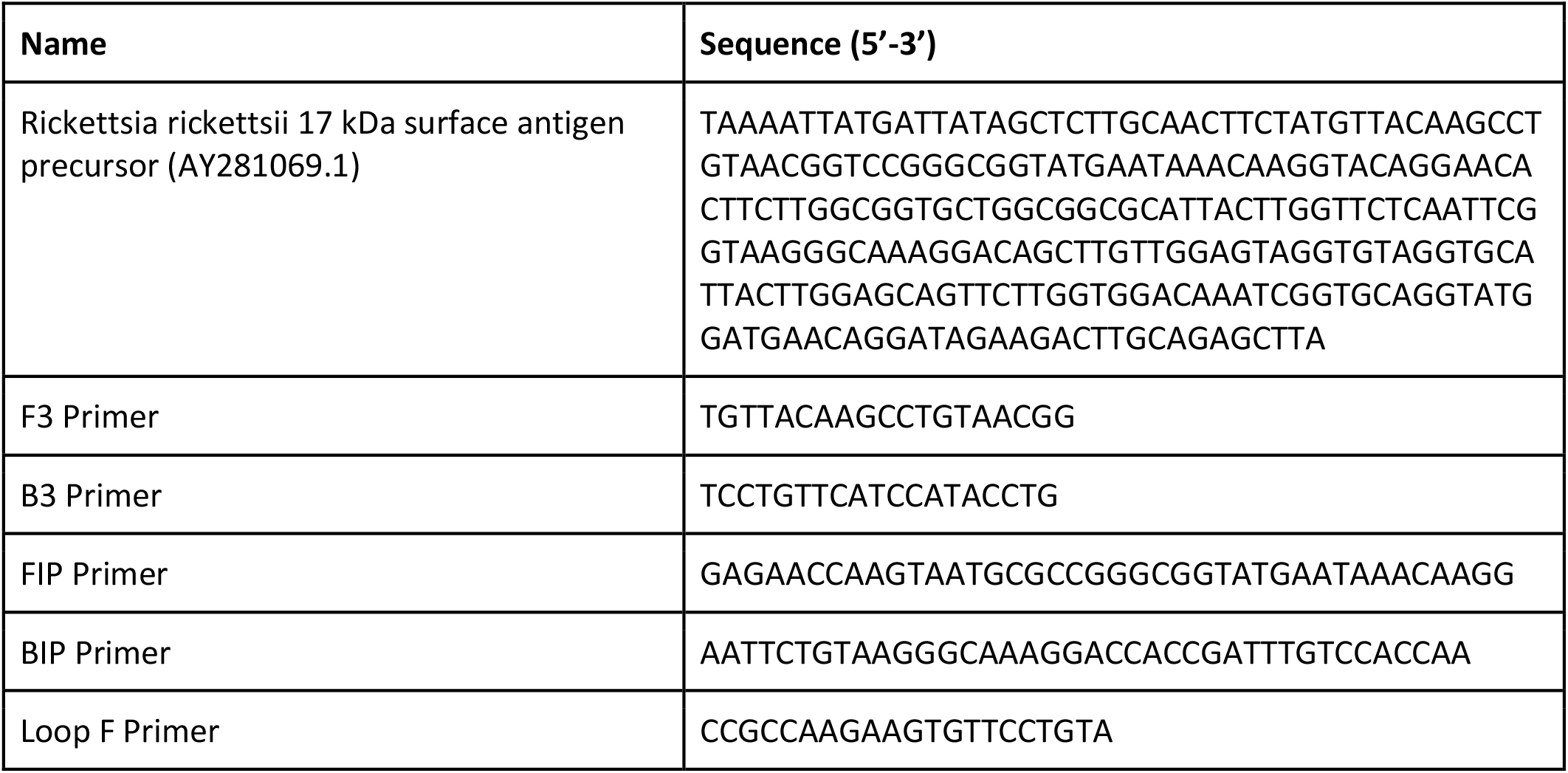
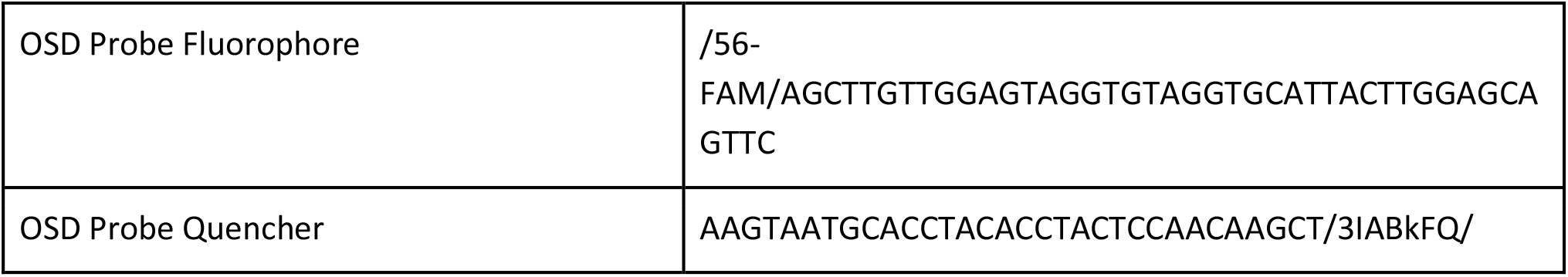
RMSF surface antigen target sequence and corresponding LAMP primer sequences for amplification.

### 3.4. Reaction Output

The LightCycler 96 created amplification curves that depict the fluorescence intensity against the number of cycles. The graphs were exported into Excel to allow for manipulation of the various chart elements. End-point assays were visualized under blue LED light with an orange filter overlay (Figure 2).

## 4. Results

The RMSF LAMP-OSD assay was tested with a serial dilution of synthetic dsDNA positive control from 10^6^ copies of DNA/μL to 10^2^ copies of DNA/μL. Successful amplification was visualized through fluorescence by the 45-minute mark for all replicates of all dilutions of the positive control samples (Figure 1, S4). Fluorescence can be clearly visualized with the naked eye by .15 fluorescence intensity. The 10^6^ copies of DNA/μL began amplification first and each ten-fold decrease in concentration resulted in a delay in both start and rate of amplification. By decreasing the concentration ten-fold from 10^6^ copies of DNA/μL down to 10^2^ copies of DNA/μL, a limit of detection was observed at 10^2^ copies of DNA/μL. None of the negative samples amplified over the hour and a half period.

**Figure 1:**
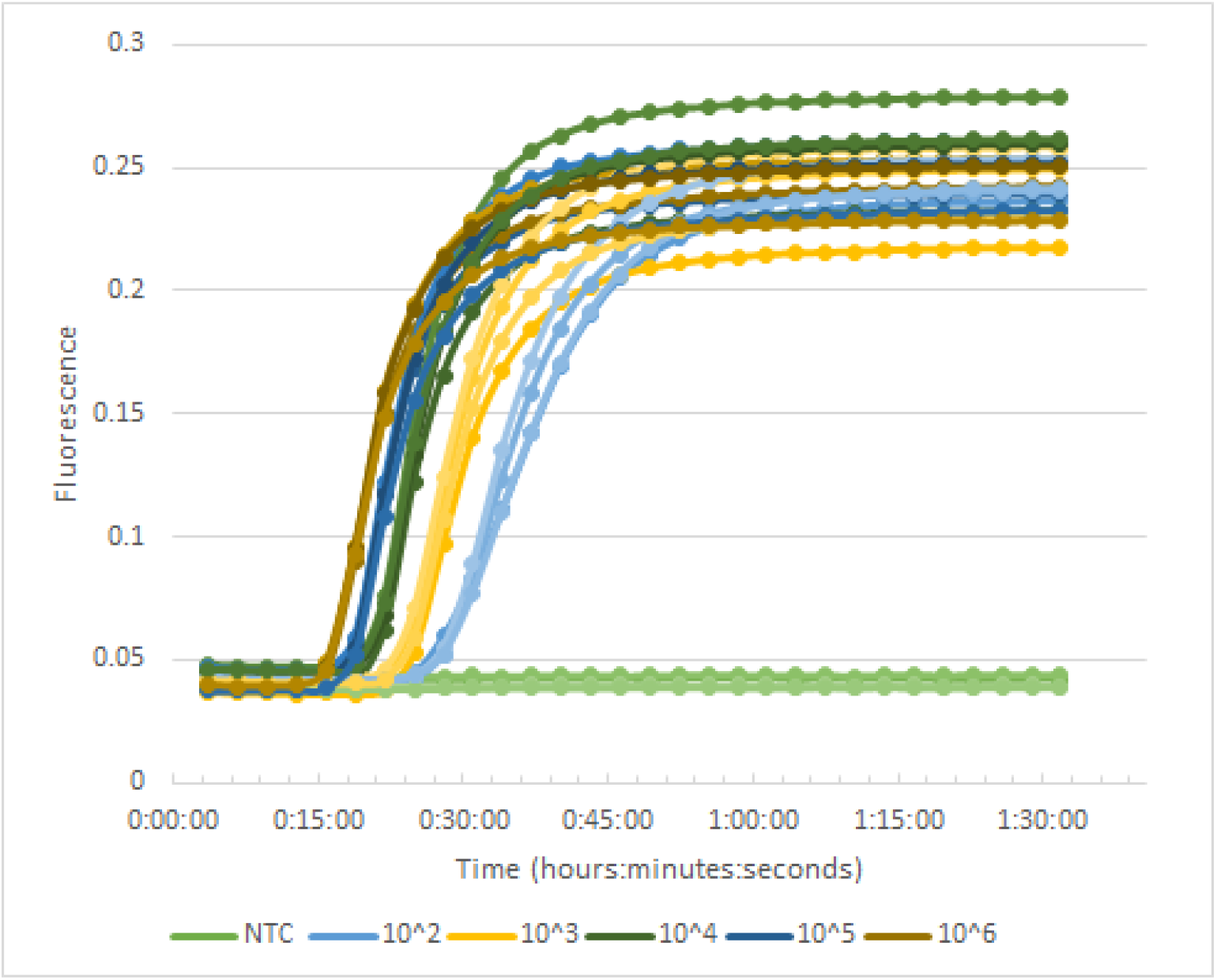
Real time OSD fluorescence curve used to determine the limit of detection. Fluorescence levels were recorded every 3 minutes and plotted against time. Successful amplification is demonstrated from 10^6^ copies of DNA/μL and serial dilutions by 10 down to 10^2^ copies of DNA/μL.

**Figure 2:**
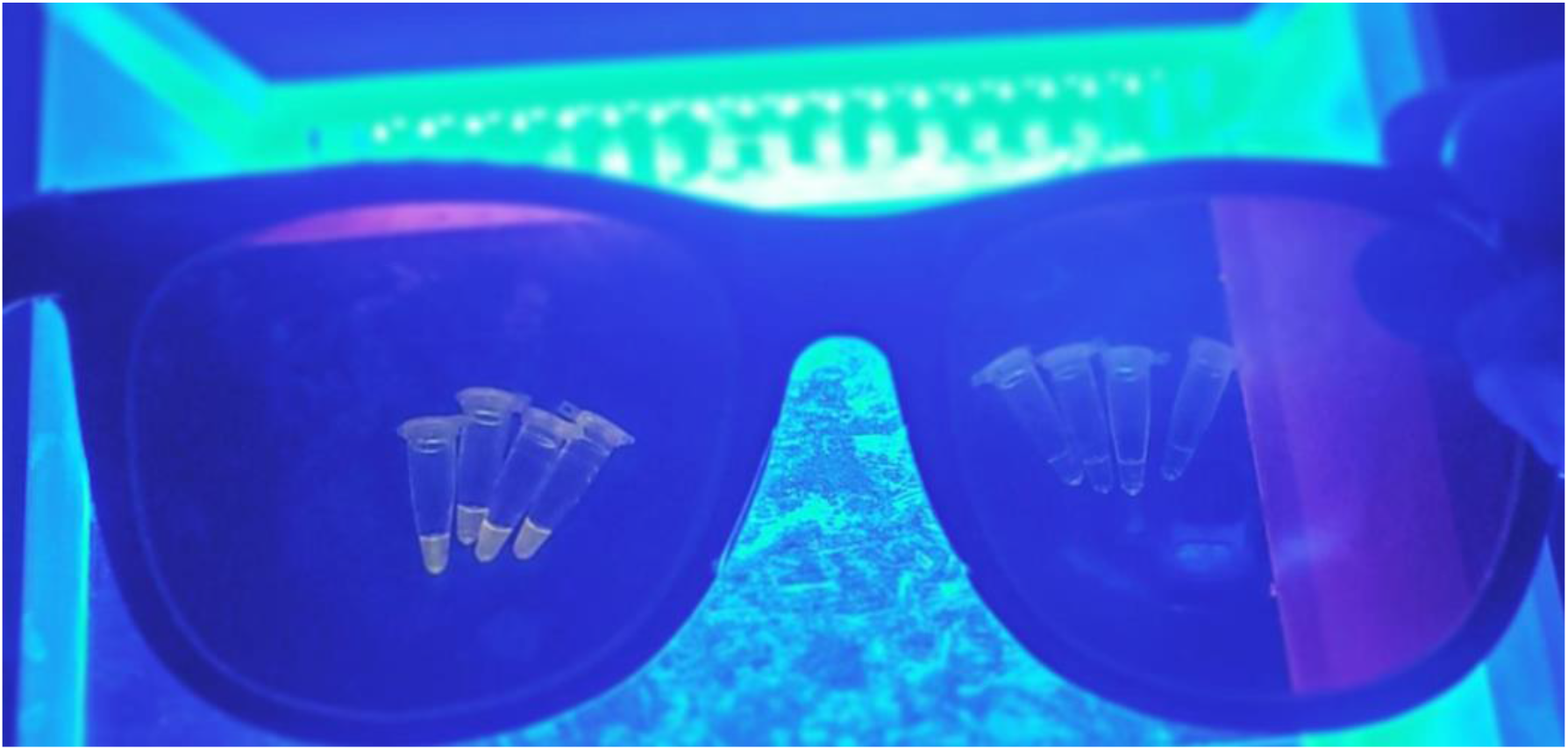
Eight end-point assays visualized under blue LEDs with orange filtered glasses. Four 10^6^ copies of DNA/μL are seen through the left lens with a clear bright yellow fluorescence. Four negative controls are seen through the right lens with non-fluorescent liquid after an hour and a half of amplification.

The LAMP-OSD protocol was then followed with a hundred-fold decrease in the concentrations of the positive samples to detect if the limit of detection would appear at an even lower rate. The data from Figure 3 supports that 10^2^ copies of DNA/μL is the limit of detection for this assay, as lower levels of DNA did not result in visualizable amplification, as the LightCycler detects fluorescence levels. Of the twelve samples with a concentration of 10^2^ copies of DNA/μL or greater, only one failed to amplify in a timely manner, while all the negative controls successfully failed to amplify.

**Figure 3:**
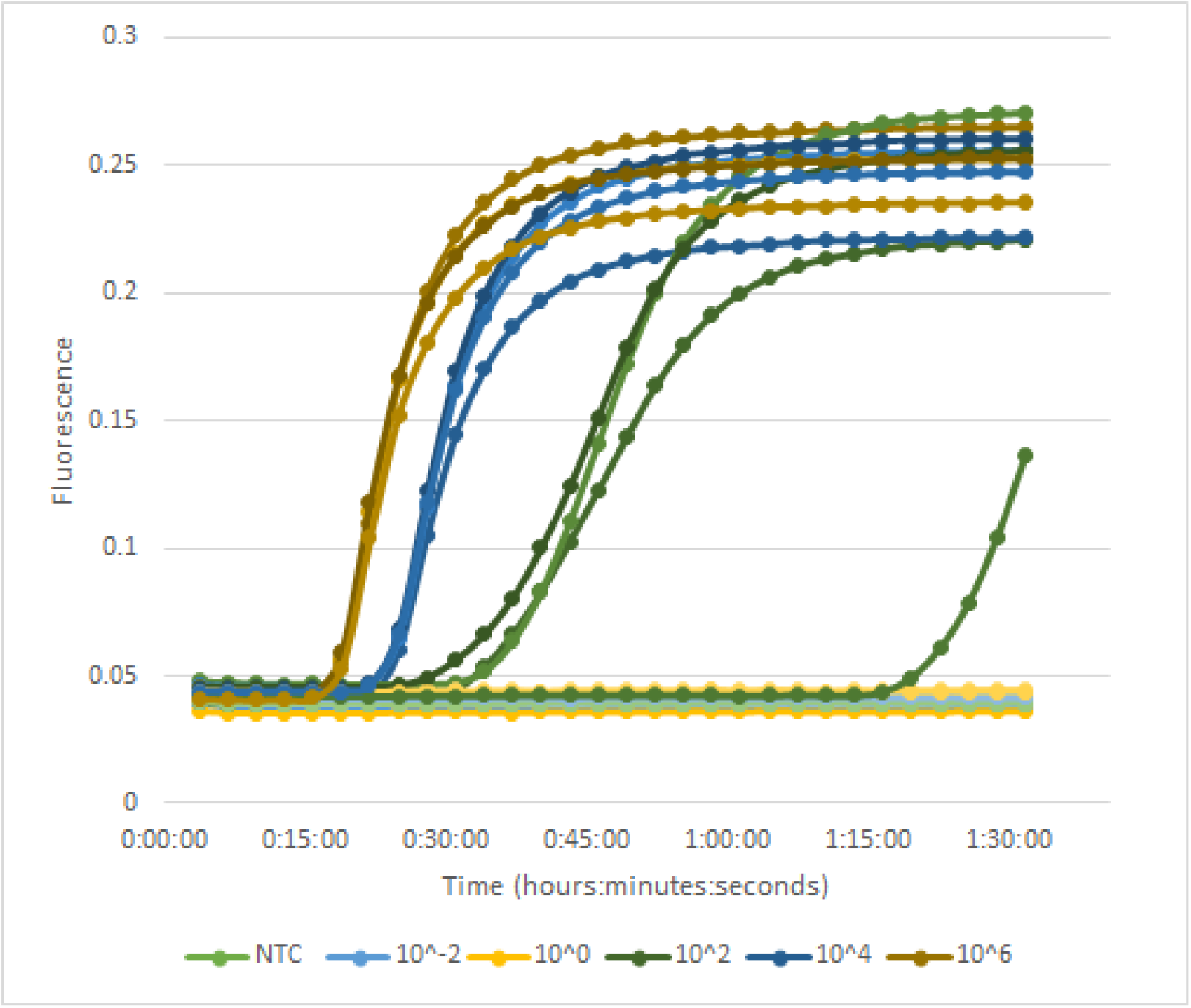
Real time OSD fluorescence curve used to determine the limit of detection. Fluorescence levels were recorded every 3 minutes and plotted against time. Successful amplification is demonstrated from 10^6^ copies of DNA/μL and serial dilutions by 100 down to 10^2^ copies of DNA/μL. Samples from 10^2^ copies of DNA/μL to 10^−2^ copies of DNA/μL demonstrate negligible to no amplification.

The OSD probe region of the R. rickettsii assay was cross-checked with the templates of various other spotted fever group diseases known for having the same treatment and having successfully been tested by Noden et al (Table 2). The designed probe shows no mutations for those species with identities above 97%, but does display mutations within the probe region for those strains more distantly related, with less than 90% strain similarity.

**Table 2:** Specificity of OSD probe versus other strains of spotted fever. The OSD probe sequence was compared with BLAST algorithm to various Rickettsial strains. (S1-S4). A consistent expect value of 0.0 was obtained for all tested strains, indicating a high significance of the match, and low background noise. Sequence identities of greater than 97% demonstrated amplification in the Noden trials.

| Strain | Expect Value | Sequence Identities | Sequence Gaps | Noden Amplification? |
| --- | --- | --- | --- | --- |
| *R. rickettsii* | 0.0 | 99% | 0% | Yes |
| *R. conorii* | 0.0 | 99% | 0% | Yes |
| *R. amblyommatis* | 0.0 | 98% | 0% | Yes |
| *R. parkeri* | 0.0 | 98% | 1% | Yes |
| *R. africae* | 0.0 | 97% | 2% | Yes |
| *R. typhi* | 0.0 | 89% | 1% | None |

## 5. Discussion/Conclusion

Spotted fever diseases have often gone undetected and untreated until the late stages due to their initially low rickettsiae levels in human fluids (Richard 2012, Biggs et al. 2016). This creates a need for a new way of testing that is more sensitive than currently used PCR counterparts. LAMP has been proven to be more sensitive in testing for Rickettsia (Noden et al. 2018) and as such, is likely to create fewer false negatives and be more reliable. Based on the findings of this research, LAMP-OSD can be used to successfully detect spotted fever group diseases SSdown to 100 copies of DNA/μL, without creating false positives, within mere hours. The previous limit of detection for synthetic DNA had been found to be .01 ng/μL using PCR. and .00001 ng/μL using SFGR-LAMP (Noden et al. 2018), however using LAMP-OSD it has been reduced down to .0000001 ng/μL. As opposed to the current confirmatory tests used by clinicians, this test can be implemented in hospital and clinic settings to return results to physicians in a timely manner to support their diagnoses. Future testing may want to test other spotted fever strains firsthand to verify their individual limits of detection, as that expectation is based on the amplification seen using SFRG-LAMP (Noden et al. 2018) it was determined that strains similar to R. *rickettsii* would behave in a relatively similar manner.

Along with the increased sensitivity, LAMP is known for being beneficial in that it is an isothermal technique and does not require complex machinery to undergo thermocycling and can produce simple yes/no readout results visualizable by the naked eye (Jiang et al. 2015, Paris et al. 2016). Given these qualities, LAMP is a much more affordable technology than PCR, needing less run time, less equipment, and less training to execute and interpret the results. As such, it would be a far more sustainable option in low-resource countries and rural areas than the current standard of PCR (Paris et al. 2016). This also opens up the possibility of large-scale vector-borne monitoring in tick populations as LAMP is a much more field-usable technology than PCR.

Future applications of this work involve partnering with local health clinics to attain comparative data on the validity of the LAMP-OSD test versus immunoserology or PCR data and to get firsthand user feedback to better enhance the tests for ease of use. Ideally, this technology can be housed in a diagnostic kit that is workable by non-experts for at-home use, especially in regions with limited access to healthcare.

## 6. Acknowledgements

We would like to acknowledge Nicholas Tran, Rithi Mulgaoker, Naomi Sequeria, and Shiv Patel for their efforts in preparing and running preliminary experiments, as well as Bob and Cathy O’Rear and The University of Texas at Austin’s Freshman Research Initiative for helping fund this work.

## Supplementary Materials

**S1:**
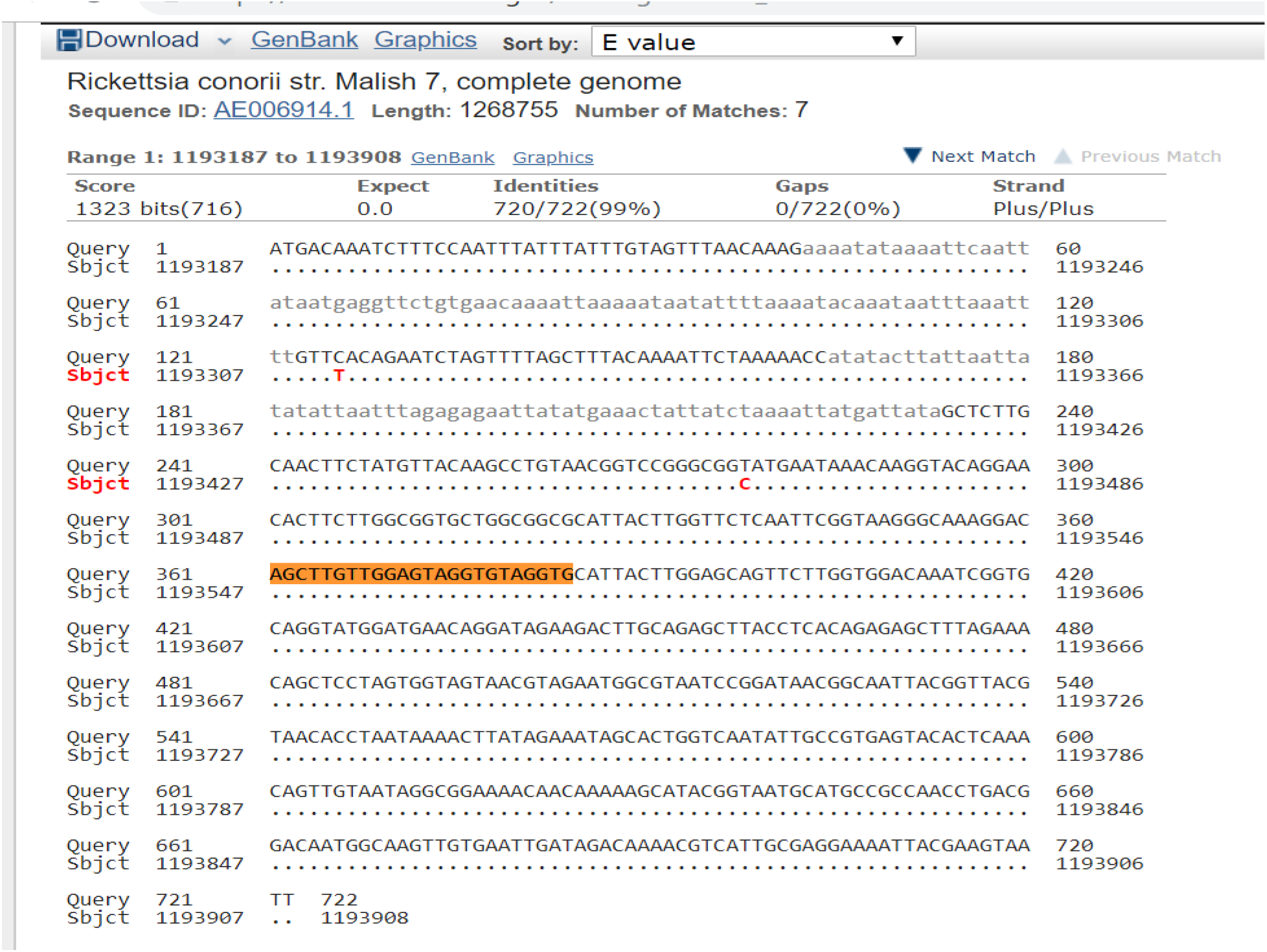
R. *rickettsii* compared with R. *conorii* and unmutated probe region identified and highlighted

**S2:**
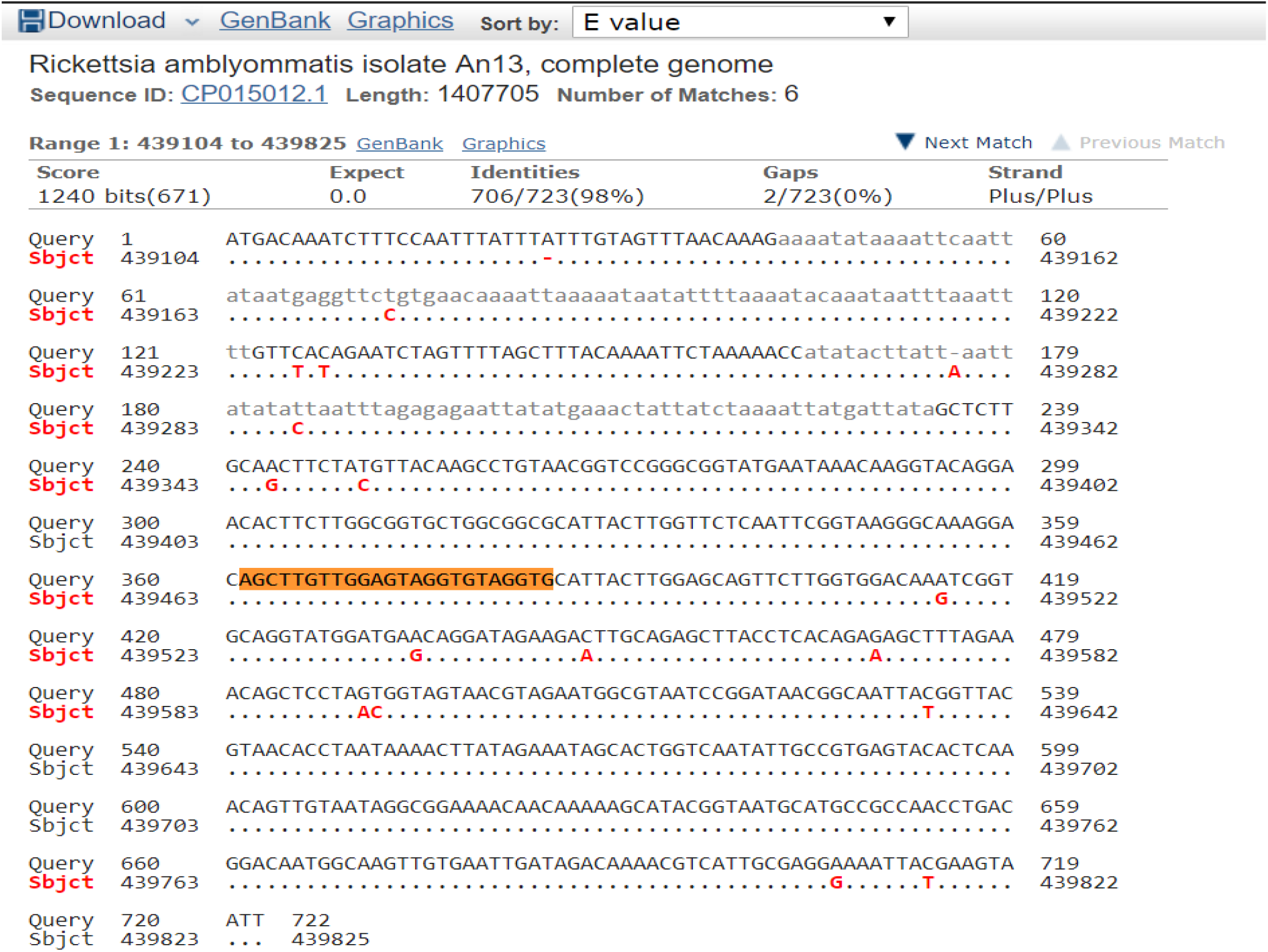
R. *rickettsii* compared with R. *amblyommatis* and unmutated probe region identified and highlighted

**S3:**
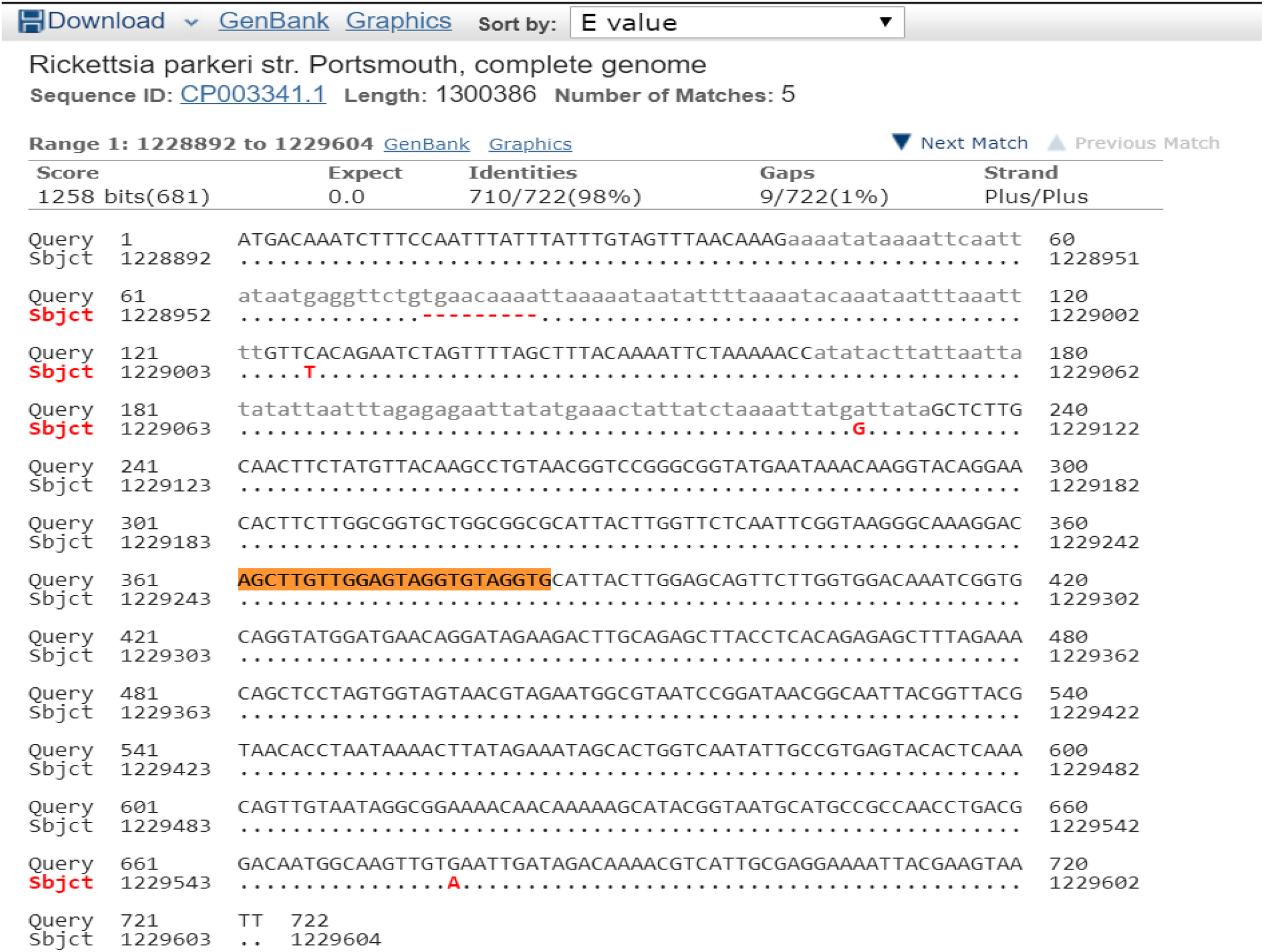
R. *rickettsii* compared with R. *parkeri* and unmutated probe region identified and highlighted

**S4:**
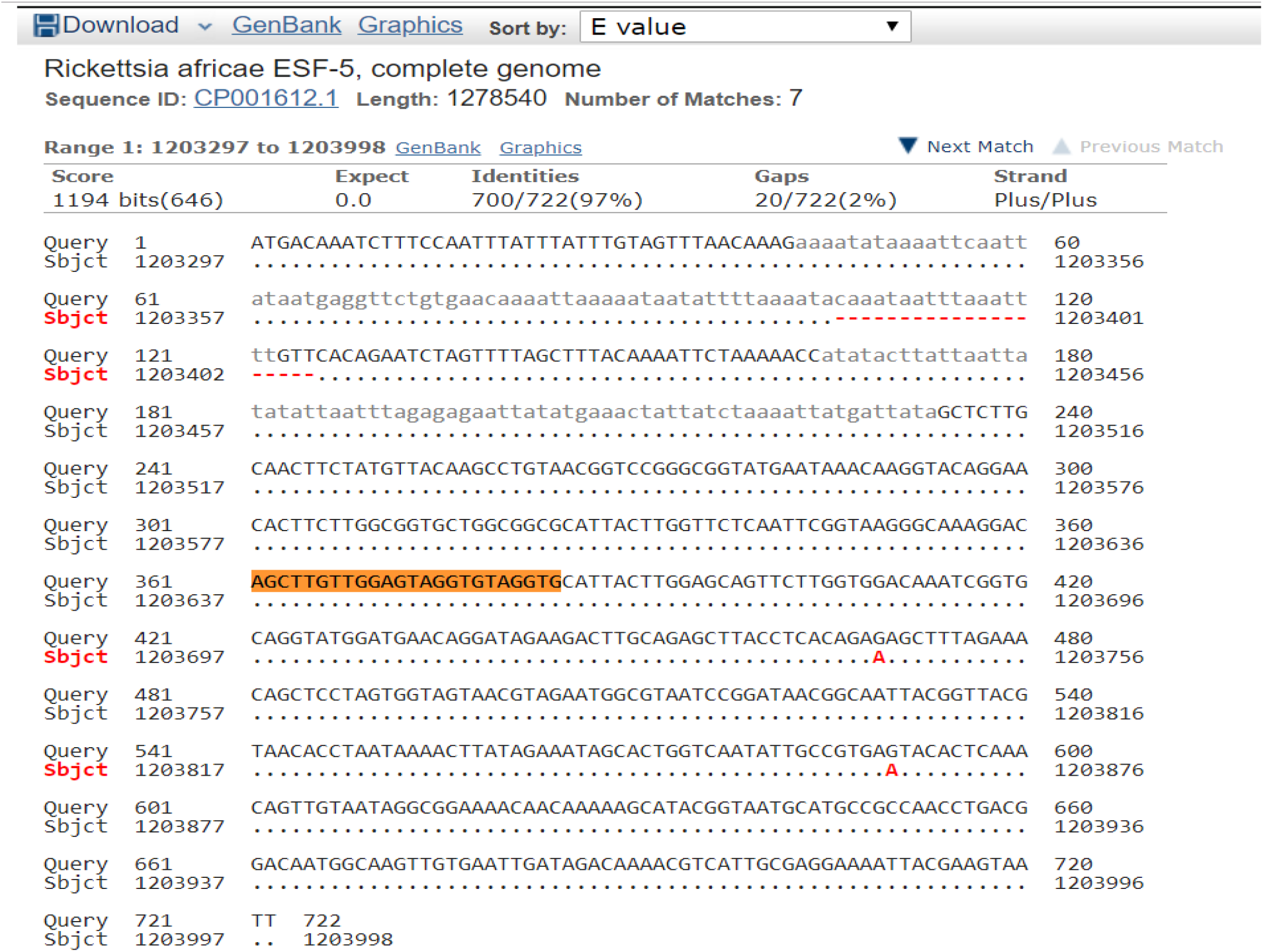
R. *rickettsii* compared with R. *africae* and unmutated probe region identified and highlighted

**S5:**
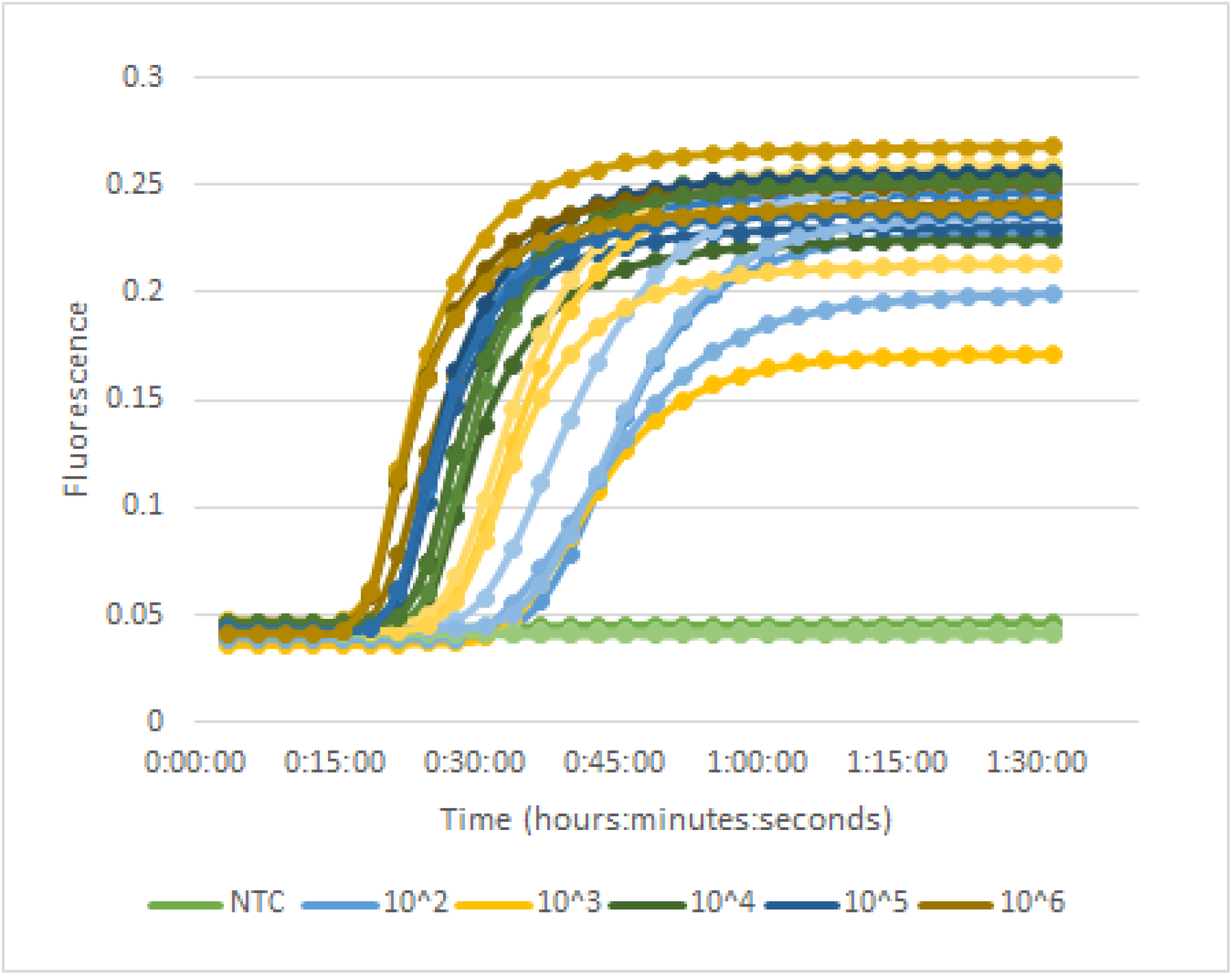
Additional limit of detection curves showing amplification of up to 10^2^ copies of DNA/μL

## References

1. Noden BH, Martin J, Carrillo Y, Talley JL, Ochoa-Corona FM. Development of a loop-mediated isothermal amplification (LAMP) assay for rapid screening of ticks and fleas for spotted fever group rickettsia. PLOS ONE. 2018 Feb 1;13(2):e0192331.

2. Richards AL. Worldwide detection and identification of new and old rickettsiae and rickettsial diseases. FEMS Immunol Med Microbiol. 2012 Feb 1;64(1):107–10.

3. Parola P, Paddock CD, Socolovschi C, Labruna MB, Mediannikov O, Kernif T, et al. Update on Tick-Borne Rickettsioses around the World: a Geographic Approach. Clinical Microbiology Reviews. 2013 Oct 1;26(4):657–702.

4. Phan JN, Lu CR, Bender WG, Smoak RM, Zhong J. Molecular Detection and Identification of Rickettsia Species in Ixodes pacificus in California. Vector Borne Zoonotic Dis. 2011 Jul;11(7):957–61.

5. Biggs HM. Diagnosis and Management of Tickborne Rickettsial Diseases: Rocky Mountain Spotted Fever and Other Spotted Fever Group Rickettsioses, Ehrlichioses, and Anaplasmosis — United States. MMWR Recomm Rep [Internet]. 2016 [cited 2020 Mar 22];65. Available from: https://www.cdc.gov/mmwr/volumes/65/rr/rr6502a1.htm

6. Agahan AL, Torres J, Fuentes-Páez G, Martínez-Osorio H, Orduña A, Calonge M. Intraocular inflammation as the main manifestation of Rickettsia conorii infection. Clin Ophthalmol. 2011;5:1401–7.

7. Raoult D, Fournier PE, Fenollar F, Jensenius M, Prioe T, de Pina JJ, et al. Rickettsia africae, a Tick-Borne Pathogen in Travelers to Sub-Saharan Africa. New England Journal of Medicine. 2001 May 17;344(20):1504–10.

8. Tick bites: First aid -Mayo Clinic [Internet]. [cited 2020 Mar 22]. Available from: https://www.mayoclinic.org/first-aid/first-aid-tick-bites/basics/art-20056671

9. Ticks Treatment [Internet]. WebMD. [cited 2020 Mar 22]. Available from: https://www.webmd.com/first-aid/ticks-treatment

10. Jiang YS, Bhadra S, Li B, Wu YR, Milligan JN, Ellington AD. Robust Strand Exchange Reactions for the Sequence-Specific, Real-Time Detection of Nucleic Acid Amplicons. Anal Chem. 2015 Mar 17;87(6):3314–20.

11. Paris DH, Dumler JS. State of the art of diagnosis of rickettsial diseases: the use of blood specimens for diagnosis of scrub typhus, spotted fever group rickettsiosis, and murine typhus. Current Opinion in Infectious Diseases. 2016 Oct;29(5):433–439.

